# Dual function of DOT1L suppresses tumor intrinsic immunogenicity in hepatocellular carcinoma

**DOI:** 10.1101/2025.05.01.651793

**Authors:** Siyuan Xu, Ruijie Gong, Songchen Liu, Jianhua Wang, Yijie Shen, Chuxuan Peng, Qin Feng, Min Luo, Fei Lan, Jia Fan, Jiabin Cai, Xianjiang Lan

## Abstract

Immune checkpoint inhibitor (ICB) therapy for many cancers remains limited in patients’ overall response rate. Discovery and development of more effective combinatorial approaches is urgent. Here, through CRISPR/Cas9 genetic screens, we identify DOT1L as a versatile epigenetic factor that functions to suppress tumor-intrinsic immunity through a dual mechanism. Depletion of DOT1L induces the expression of transposable elements and subsequent type I interferon (IFN) response, and meanwhile lowers *ZEB1* levels to further unleash the expression of immune-related genes. In turn, we demonstrate that DOT1L loss or treatment with the clinical stage inhibitor EPZ-5676 sensitizes tumors to ICB with increased immune infiltration in mice. More importantly, EPZ-5676 treatment alone is sufficient to enhance antitumor immunity in humanized mice. TCGA data analysis reveals an inverse correlation between DOT1L expression and IFN signatures across multiple cancer types. These findings provide a rationale for targeting DOT1L to improve tumor immunogenicity and overcome immunotherapy resistance.

**One Sentence Summary:** CRISPR genetic screens identified DOT1L as a potential suppressor of tumor intrinsic immunogenicity

## INTRODUCTION

Immune checkpoint blockade (ICB) has broad impact on cancer immunotherapy, with several antibodies targeting the programmed cell death 1 (PD1)–PD1 ligand 1 (PD-L1) axis or cytotoxic T lymphocyte antigen 4 (CTLA-4) approved for use in a number of different cancers (*1, 2*). However, despite these advancements, a substantial proportion of patients do not derive clinical benefits from ICB due to tumor heterogeneity or the immunosuppressive tumor microenvironment (TME) (*3*). Therefore, identifying better biomarkers and development of novel therapeutic strategies to overcome intrinsic or adaptive resistance for immunotherapy is urgently needed.

The immune checkpoint genes, such as the innate immune regulator cluster of differentiation 47 (CD47) and the adaptive immune checkpoint PD-L1, are typically up-regulated by tumors to evade the host immune system (*4, 5*). Inhibition of the immune checkpoint genes can reshape immunosuppressive TME and thus enhance the activity of tumor-infiltrating T cells, macrophages or natural killer (NK) cells across various cancers (*6*). For example, MYC suppression enhanced the antitumor immune response through directly down-regulating both *CD47* and *PD-L1* expression (*7*). In addition, triggering the innate immune response within tumors via viral-mimicry-inducing therapies by epigenetic perturbation can remodel TME and sensitize the ICB treatment (*8, 9*). Several studies have revealed that loss of epigenetic factors, including the DNA methyltransferase DNMT1 (*10, 11*), the histone methyltransferase SETDB1 (*12*), the histone demethylases KDM1A or KDM5B (*13, 14*), and the chromatin remodeler ARID1A or PBAF (*15, 16*), can induce antitumor immunity through distinct mechanisms in melanoma or colorectal cancer. However, the epigenetic factors modulating the expression of immune checkpoints or tumor cell-intrinsic immunity in other cancer types are still largely unknown.

Here, using the immune checkpoint molecules CD47 and PD-L1 as the reporters, we carried out CRISPR genetic screens in hepatocellular carcinoma (HCC) cell line and identified disruptor of telomeric silencing 1like (DOT1L), the only histone lysine methyltransferase for histone H3 on lysine 79 (H3K79) mono-, di-, and tri-methylation (*17, 18*), as a novel repressor of tumor cell-intrinsic immunity. We demonstrate that DOT1L promotes tumor immune evasion through restraining the expression of endogenous transposon elements (TEs) and meanwhile activating *ZEB1* expression to further repress the immune-related gene levels in an enzymatic activity-dependent manner in humans. Dot1l loss or Dot1l activity inhibition leads to reversal of resistance to anti-PD-L1 therapy in murine tumors. Furthermore, DOT1L inhibitor EPZ-5676 treatment alone significantly improves the antitumor immunity in an HCC xenograft humanized mouse model. Together, these findings might open new paths for targeting epigenetic regulator DOT1L to overcome immunotherapy resistance.

## RESULTS

### CRISPR screens identify DOT1L as a novel immune checkpoint repressor

To identify novel regulators of immune checkpoint *CD47* or *PD-L1* expression, we screened a sgRNA library (table S1) targeting the chromatin regulatory domains (*19*) of most epigenetic factors (∼219 total; on average 6 sgRNAs per domain). A lentiviral vector library encoding the sgRNAs was used to infect HuH-7 cells that stably express spCas9 (HuH-7-Cas9). The top 15% and bottom 15% of CD47 or PD-L1 expressing cells were purified by anti-CD47 or PD-L1 fluorescence activated cell sorting (FACS). Since IFNγ is one of the most essential cytokines in the immune response and known to stimulate both *CD47* and *PD-L1* expression (*20, 21*), we also performed the screens in the presence of IFNγ. Subsequently, the sorted cells were subject to deep sequencing to assess the representation of each sgRNA in the two cell populations (Fig.1A).

**Fig. 1.**
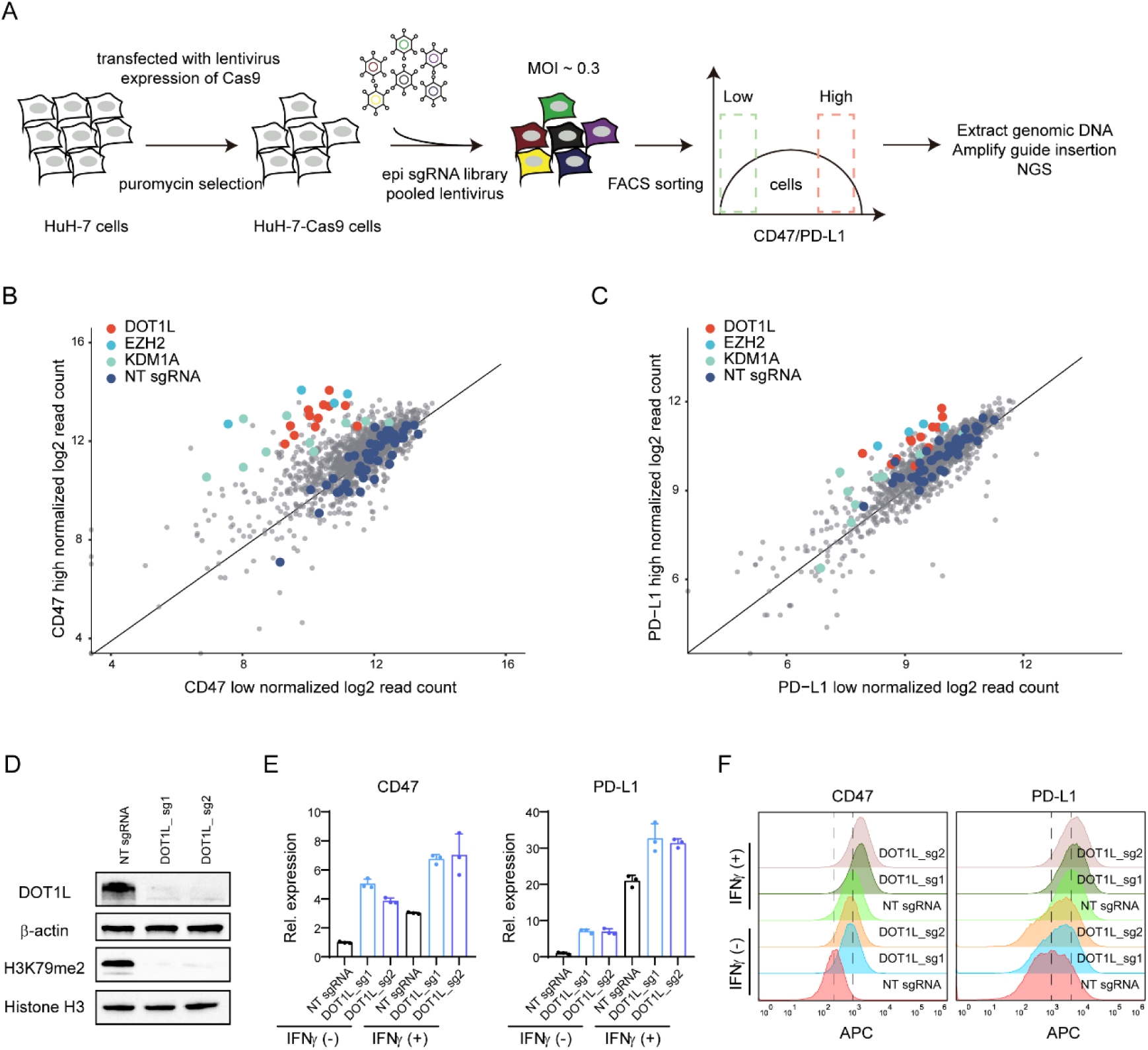
FACS-based CRISPR/Cas9 genetic screens in HuH-7 cells. (**A**) Schematic of the screening strategy. epi, epigenetic; FACS, fluorescence-activated cell sorting; NGS, Next-generation sequencing. (**B** and **C**) Scatterplot of the screen results. Control sgRNAs (dark blue) are scattered randomly across the diagonal. EZH2 (light blue) and KDM1A (green) represent positive control sgRNAs. NT, non-targeting. (**D**) Immunoblotting analysis using whole-cell lysates from HuH-7 cells transduced with the indicated sgRNAs. (**E**) *CD47* or *PD-L1* mRNA levels in HuH-7 cells upon DOT1L depletion with or without IFNγ treatment for 48 h. Results are shown as mean ± SD (n = 3). (**F**) CD47 or PD-L1 flow cytometry analyses of WT and DOT1L-depleted HuH-7 cells with or without IFNγ treatment for 48 h.

As expected, positive control sgRNAs targeting the known IFN pathways repressor genes KDM1A and EZH2 (*13, 22*) were enriched in the CD47 and PD-L1-high populations (Fig. 1, B and C). Besides, all twelve sgRNAs targeting DOT1L were significantly enriched in both CD47 and PD-L1-high cells with or without IFNγ treatment (Fig. 1, B and C, fig. S1, A to D, and table S1), suggesting that DOT1L may function as a direct or indirect repressor of *CD47* and *PD-L1* expression. To validate the screening results, six independent sgRNAs targeting DOT1L and non-targeting control sgRNA were stably introduced into HuH-7-Cas9 cells, respectively. Depletion of DOT1L strongly increased the expression levels of *CD47* and *PD-L1* without IFNγ treatment (Fig. 1, D and E, and fig. S1, E to G). Notably, the mRNA and protein levels of CD47 and PD-L1, determined by RT-qPCR and flow cytometry experiments, were further elevated for hyperactivation in DOT1L-depleted HuH-7-Cas9 cells following IFNγ stimulation, compared with the baseline (Fig. 1, E and F). Moreover, the results of DOT1L deficiency in another HCC cell line HepG2 were similar to those in HuH-7 cells (fig. S1, H to J). Collectively, these findings demonstrate that DOT1L acts as a novel repressor of dual immune checkpoint *CD47* and *PD-L1*.

### Loss of DOT1L triggers a cell-intrinsic immune response and primes response to IFNγ

Previous studies have shown that activation of innate immune response can lead to up-regulation of the immune checkpoint expression (*13, 22, 23*). To further determine how DOT1L regulates *CD47* and *PD-L1*, we performed RNA-sequencing (RNA-seq) experiments in HuH-7 cells and found that the number of up-regulated genes was far more than that of down-regulated genes in both independent DOT1L sgRNAs (Fig. 2, A and B, fig. S2A, and table S2 and S3), suggesting that DOT1L acts as a transcriptional repressor at most targets. GO enrichment analysis showed that gene sets related to IFN response and antiviral response were significantly enriched among up-regulated genes in DOT1L-depleted cells, compared with WT cells, and ranked at the top (Fig. 2C). IFN signatures and immune signaling pathways were also among the top five most significantly enriched gene sets in DOT1L-deficient cells according to gene set enrichment analysis (GSEA; Fig. 2, D and E, and fig. S2B). Moreover, compared with the baseline, up-regulation of these pathways in DOT1L knockout (KO) cells with IFNγ treatment was confirmed by RNA-seq, followed with GSEA, GO enrichment analysis, and KEGG pathway analysis (fig. S2, C to F). Furthermore, the up-regulated expression of IFN stimulated genes (ISGs) encoding antiviral responses (*OAS1* and *OAS3*), pro-inflammatory cytokines (*CXCL10* and *CXCL11*), and antigen presentation genes (*HLA-A*, *B*, and *C*) in DOT1L deficient cells with or without IFNγ treatment were validated by RT-qPCR experiments (fig. S2, G and H), consistent with the RNA-seq results (Fig. 2, F and G). Additionally, major histocompatibility complex class I (MHC I) cell surface expression was also elevated at baseline and hyper-induced following IFNγ stimulation in DOT1L deficient cells via flow cytometry (Fig. 2H). In line with HuH-7 cells, depletion of DOT1L in HepG2 cells led to the up-regulation of ISGs with or without IFNγ treatment (fig. S2, I and J).

**Fig. 2.**
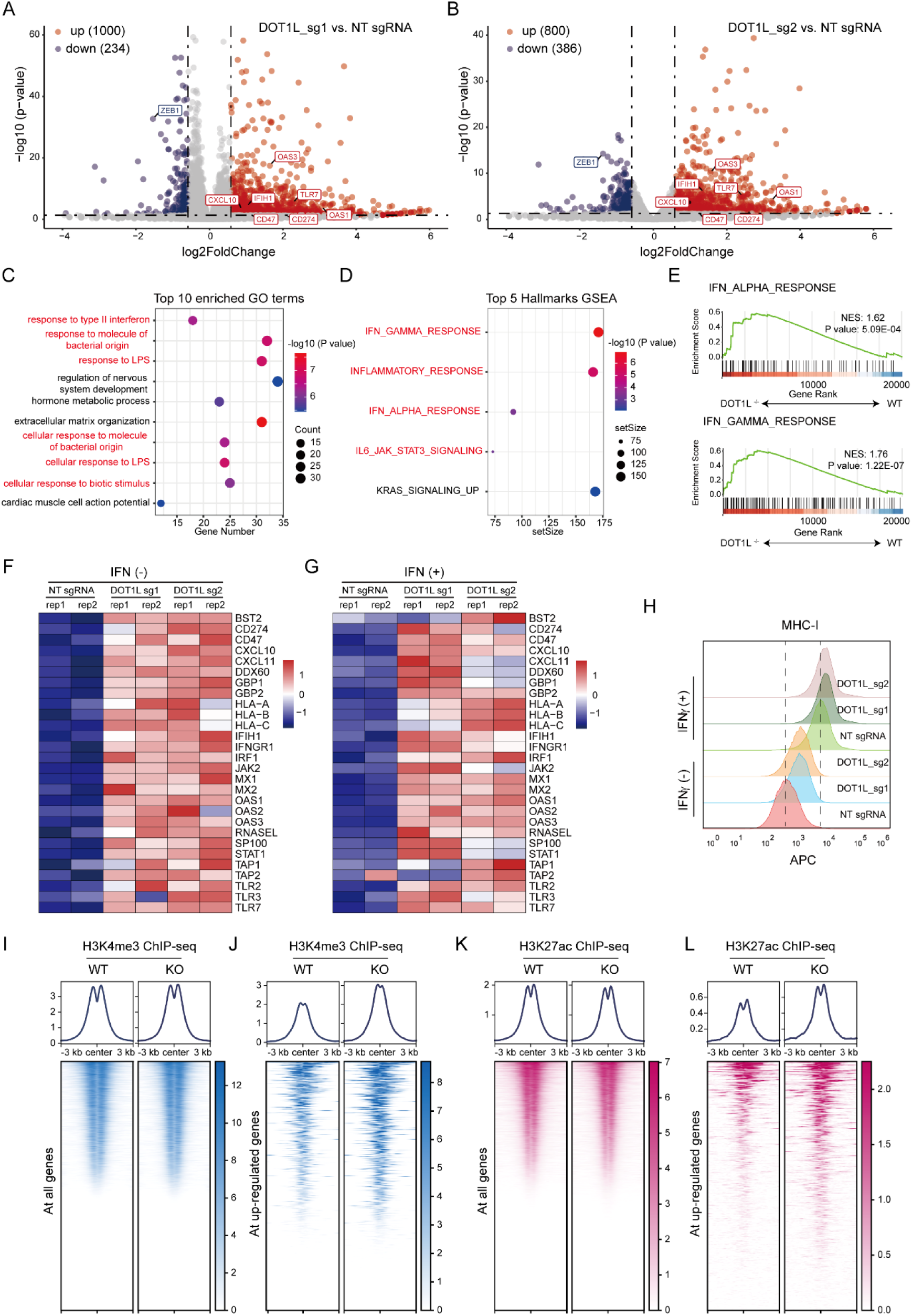
DOT1L negatively regulates interferon signaling pathway. (**A** and **B**) Volcano plot of RNA-seq analysis in WT and DOT1L-depleted HuH-7 cells. NT, non-targeting. (**C**) Gene ontology (GO) analysis of RNA-seq data showing top 10 pathways that are up-regulated upon DOT1L depletion in HuH-7 cells. (**D**) Gene Set Enrichment Analysis (GSEA) of top 5 hallmark gene sets enriched in DOT1L-depleted HuH-7 cells. (**E**) GSEA of RNA-seq data showing IFNα (top) and IFN**γ** (bottom) gene sets that are up-regulated upon DOT1L depletion in HuH-7 cells. (**F** and **G**) Heatmaps of selected immune-related genes whose expression is higher in DOT1L-depleted HuH-7 cells with or without IFNγ treatment for 48 h. (**H**) Flow cytometry histograms of MHC I in sgDOT1L HuH-7 cells with or without IFNγ treatment for 48 h. (**I** and **J**) Binding density heatmap showing H3K4me3 distribution at the promoters of all genes (I) or the up-regulated genes (J) caused by DOT1L KO in HuH-7 cells. (**K** and **L**) Binding density heatmap showing H3K27ac distribution at the promoters of all genes (K) or the up-regulated genes (L) caused by DOT1L KO in HuH-7 cells.

Epigenetic modifications play important roles in gene regulation. We therefore reasoned that loss of DOT1L might alter histone modifications of those up-regulated immune-related genes. To this end, we performed chromatin immunoprecipitation with high-throughput sequencing (ChIP-seq) experiments against histone 3 trimethylation on lysine 4 (H3K4me3) and histone 3 acetylation on lysine 27 (H3K27ac), linked to active transcription. We found that the distributions of H3K4me3 and H3K27ac in all gene promoters were unaffected by DOT1L depletion in HuH-7 cells without IFNγ treatment (Fig. 2, I and K). However, sequence motif analysis of significantly increased H3K4me3 peaks showed enrichment of binding sites of IRF family and IFN-stimulated response element (ISRE; fig. S2K). Additionally, we found enrichment of NF-κB, AP-1, NFAT, STAT3 and IRF1 binding motifs in chromatin regions that gained H3K27ac peaks after DOT1L loss (fig. S2L). These observations support the activation of IFN signaling pathways in DOT1L depleted HuH-7 cells. Moreover, when only the up-regulated genes were included for comparison, the signal density of both H3K4me3 and H3K27ac were remarkably increased (Fig. 2, J and L), suggesting that the H3K4me3 and H3K27ac changes on immune-related genes are positively correlated with its gene expression. Together, these data demonstrate that DOT1L deficiency activates the cell-intrinsic immune response and enhances response to IFNγ.

### The DOT1L methyltransferase activity is vital to restrain cell-intrinsic immune response

To further study whether the DOT1L methyltransferase activity is required for repressing the IFN pathway activation, we utilized the potent DOT1L inhibitor EPZ-5676 (*24*) in cells. Treatment of HuH-7 cells with EPZ-5676 at increasing concentrations decreased the global levels of histone H3K79me2 mark, without affecting the DOT1L protein levels (Fig. 3A). Strikingly, the mRNA levels and cell surface protein levels of CD47 and PD-L1 were elevated in a dose-dependent manner with EPZ-5676 treatment (Fig. 3, B and C), recapitulating the effect of DOT1L-depleted HuH-7 cells (Fig. 1, E and F). To further understand the transcriptome changes after EPZ-5676 treatment, we carried out RNA-seq experiments and again found that the number of up-regulated genes was much more than that of down-regulated genes (Fig. 3D and table S4). GO enrichment analysis revealed that the top terms among up-regulated genes were related to immune response with EPZ-5676 treatment, compared with the DMSO control (Fig. 3E). Some of the up-regulated immune-related genes were further validated by RT-qPCR experiments (Fig. 3, F and G). We also confirmed that the MHC I cell surface expression was elevated via flow cytometry (Fig. 3H). Moreover, GSEA revealed the up-regulation of IFN-dependent and cell-intrinsic immune signaling pathways by EPZ-5676 treatment (Fig. 3I and fig. S3A). Notably, the commonly up-regulated 366 genes in both DOT1L depleted and EPZ-5676 treated cells were associated with IFN pathway activation (fig. S3, B and C). Additionally, EPZ-5676 treatment enhanced the response of immune-related genes to IFNγ (fig. S3, D to G), consistent with DOT1L-deficient cells. Again, in line with that in HuH-7 cells, treatment of HepG2 cells by EPZ-5676 led to activation of IFN signaling pathways (fig. S3, H to K).

**Fig. 3.**
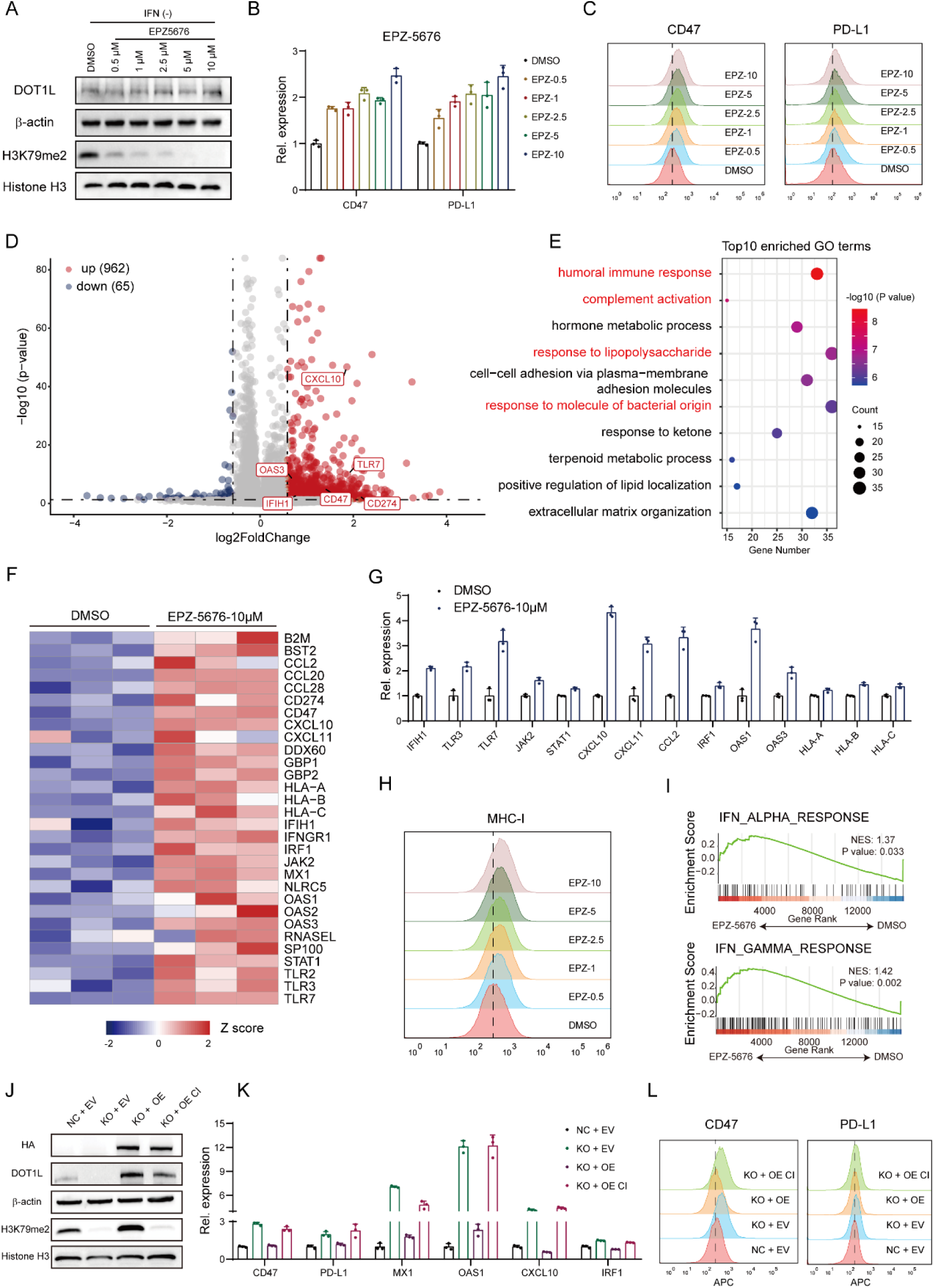
The DOT1L methyltransferase activity is vital to repress interferon signaling pathway. (**A**) Immunoblotting analysis using whole-cell lysates from HuH-7 cells with EPZ-5676 treatment for 7 days. (**B**) RT-qPCR analysis of *CD47* and *PD-L1* mRNA levels of HuH-7 cells with EPZ-5676 treatment for 7 days. Results are shown as mean ± SD (n=3). (**C**) Flow cytometry histograms of CD47 and PD-L1 in HuH-7 cells with EPZ-5676 treatment for 7 days. (**D**) Volcano plot of RNA-seq analysis in DMSO and EPZ-5676 treatment HuH-7 cells. (**E**) GO analysis of RNA-seq data showing top 10 pathways that are up-regulated in HuH-7 cells with EPZ-5676 treatment. (**F**) Heatmaps of selected immune-related genes whose expression is higher in EPZ-5676 treatment for 7 days. (**G**) Expression levels of immune-related genes by RT-qPCR analysis. Results are shown as mean ± SD (n=3). (**H**) Flow cytometry histograms of MHC I in HuH-7 cells with EPZ-5676 treatment for 7 days. (**I**) GSEA of RNA-seq data showing IFNα (top) and IFNγ (bottom) gene sets that are up-regulated upon DOT1L depletion in HuH-7 cells. (**J**) Immunoblotting analysis of the DOT1L expression after overexpression of WT or catalytic inactive (CI) HA-DOT1L in DOT1L-depleted HuH-7 cells. (**K**) RT-qPCR analysis of immune-related genes after rescue of WT or CI HA-DOT1L in DOT1L-depleted HuH-7 cells. Results are shown as mean ± SD (n=3). (**L**) Flow cytometry histograms of CD47 and PD-L1 after overexpression of WT or CI HA-DOT1L in DOT1L-depleted HuH-7 cells.

Meanwhile, we performed rescue experiments through restoring the HA-tagged WT or catalytic inactive (CI) form of DOT1L in DOT1L deficient HuH-7 cells. Immunoblot analysis showed comparable protein levels between WT and CI DOT1L (Fig. 3J). Notably, the transcription levels of immune-related genes were remarkably restored by WT DOT1L, rather than the catalytic mutant (Fig. 3K). The cell surface levels of CD47 and PD-L1, as determined by flow cytometry, were also rescued by WT DOT1L (Fig. 3L). Taken together, these findings indicate that the DOT1L enzymatic activity plays a crucial role in repressing the cell-intrinsic immune response.

### DOT1L represses retroelement expression to restrain type I IFN response

Previous studies have reported that depletion of histone methyltransferase SETDB1 and EZH2, or demethylase KDM1A and KDM5B activates retroelement expression and thus induces an IFN response via MDA5/RIG-1 dsRNA pathway and/or cGAS/STING pathway (*12–14, 22*). To understand the effect on TEs upon DOT1L deficiency in HuH-7 cells, we carried out total RNA-seq experiments and found that 440 TEs were significantly up-regulated and only 6 TEs were down-regulated (Fig. 4A), mainly including long interspersed nuclear elements (LINEs), long terminal repeat (LTR), short interspersed nuclear elements (SINEs) and DNA transposon (Fig. 4B and fig. S4, A and B). Up-regulation of the retroelement expression was further validated by RT-qPCR analysis in DOT1L-depleted and EPZ-5676 treated HuH-7 cells (Fig. 4C and fig. S4C). Since DOT1L is an epigenetic modifier associated with chromatin regulation, we next performed assay for transposase-accessible chromatin with high-throughput sequencing (ATAC-seq) to examine the chromatin landscape and found that DOT1L loss had minimal effect on global chromatin accessibility (fig. S4D), while increased the chromatin accessibility on TEs (Fig. 4D). Consistently, these more accessible TE regions were deposited with increased H3K4me3 and H3K27ac marks upon DOT1L loss (Fig. 4E). Moreover, the fraction of TEs (∼38%) in more accessible sites was higher than that of TEs (∼22%) in less accessible sites after DOT1L depletion (fig. S4, E and F), in line with the total RNA-seq data.

**Fig. 4.**
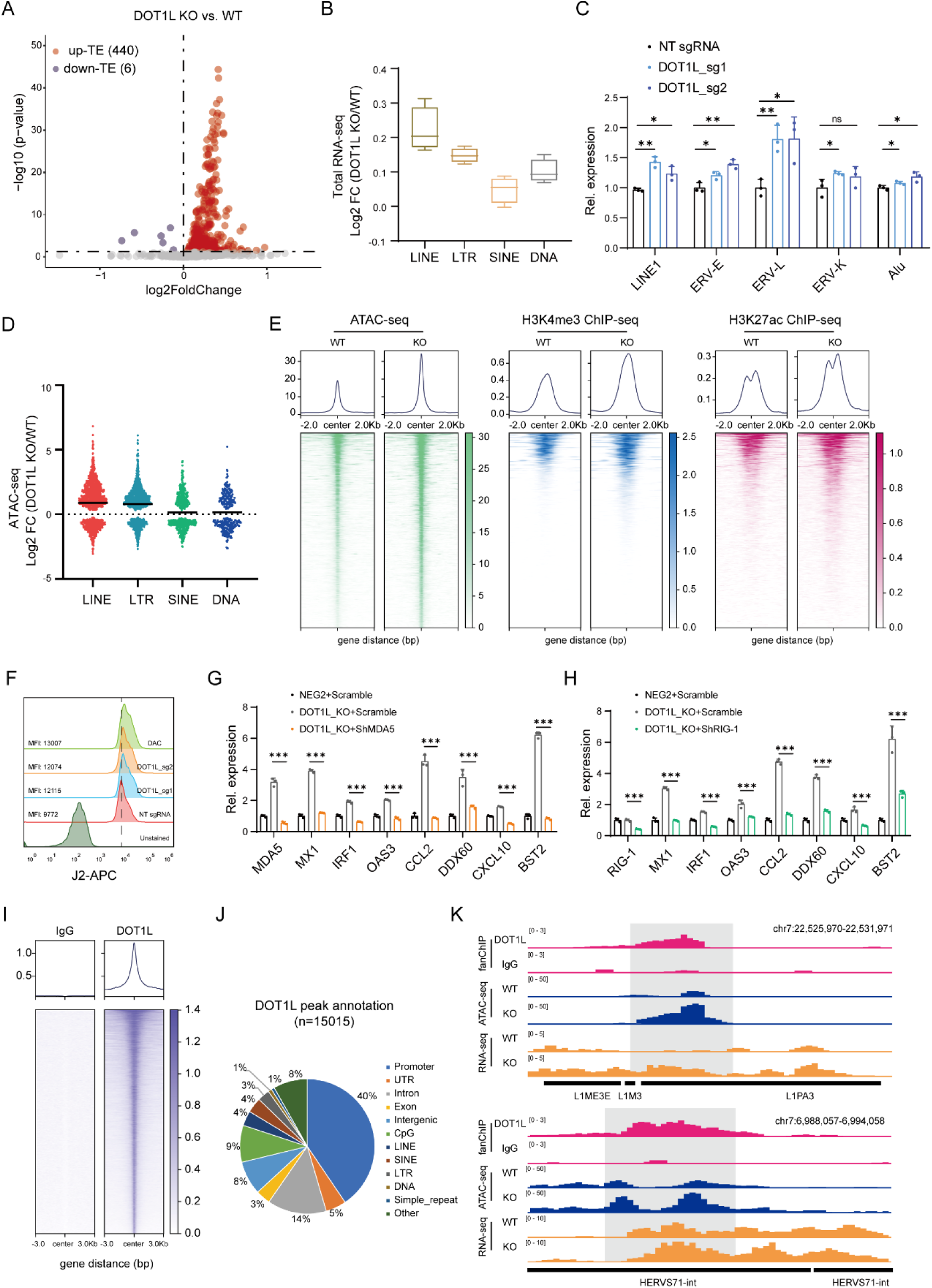
DOT1L restrains intracellular dsRNA formation. (**A**) Volcano plot shows transcriptome analysis of transposon elements (TEs) expression after DOT1L KO in HuH-7 cells. (**B**) The expression changes of four different categories of TEs in WT and DOT1L KO HuH-7 cells. (**C**) RT-qPCR analyses of TEs in WT and DOT1L-depleted HuH-7 cells. Results are shown as mean ± SD (n=3). n.s., not significant, \**P* < 0.05, \*\**P* < 0.01, unpaired Student’s *t*-test. (**D**) The ATAC-seq signal changes of four different categories of TEs in WT and DOT1L KO HuH-7 cells. (**E**) Binding density heatmap of ATAC-seq, H3K4me3, and H3K27ac ChIP-seq signals of up-regulated TEs caused by DOT1L KO in HuH-7 cells. (**F**) Flow cytometry histograms of dsRNA staining of WT or DOT1L KO HuH-7 cells. DAC, Decitabine served as a positive control. (**G** and **H**) RT-qPCR analysis of immune-related genes in DOT1L-depleted HuH-7 cells transduced with shRNA against scramble, MDA5 (G), or RIG-I (H). Results are shown as mean ± SD (n=3). \*\*\**P* < 0.001, unpaired Student’s *t*-test. (**I**) Heatmaps of IgG and DOT1L signals by fanChIP in HuH-7 cells. (**J**) Genomic annotations of DOT1L binding peaks by fanChIP in HuH-7 cells. (**K**) DOT1L fanChIP signals, ATAC-seq, and RNA-seq profiles in HuH-7 cells of LINEs (top) or LTR (bottom).

Both sense and antisense transcription of retroelements may lead to the generation of immunogenic double-strand RNA (dsRNA), further provoking the type I IFN pathway (*9*). Notably, depletion of DOT1L in HuH-7 cells led to accumulation of dsRNA, as determined by flow cytometry with J2 antibody (Fig. 4F). Importantly, abrogation of the intracellular dsRNA sensors (MDA5 or RIG-1) significantly diminished the induction of ISGs in DOT1L-depleted cells (Fig. 4, G and H). Furthermore, knockdown of type I IFN receptor (IFNAR1), but not type II IFN receptor (IFNGR1), efficiently restored the repression of ISGs (fig. S4, G and H). These data indicate that the RNA-sensing pathways contribute to the activation of the type I IFN pathway that is induced by DOT1L depletion.

DOT1L is usually associated with transcription activation via its histone substrate H3K79me2 (*25, 26*), but it also participates in transcription repression (*27, 28*). To determine how DOT1L regulates TE expression, we performed fanChIP experiments (*29*) in HuH-7 cells. Notably, among 15015 high-confidence DOT1L binding peaks, 12% located at TEs (Fig. 4, I and J). Of note, the majority of DOT1L peaks bound to the gene promoters (∼40%; Fig. 4J), consistent with that DOT1L overlaps with RNAP II at actively transcribed genes (*26*). Importantly, given that DOT1L depletion mainly enhanced the transcription and the chromatin accessibility of LINEs and LTR (Fig. 4, B and D), we discovered that a fraction of these LINEs and LTR (12.3%) were bound by DOT1L (Fig. 4K and fig. S4, I and J), suggesting its transcription repressive role. Moreover, Cistrome-GO analysis (*30*) of the ATAC-seq data for the DOT1L-bound LINEs and LTR revealed that genes with increased regulatory potential in DOT1L deficient cells showed significant enrichment of IFN signaling pathways (fig. S4, K and L). However, these regions were enriched with little inert H3K79me2 signals (fig. S4M), implying that DOT1L appears to repress these LINEs and LTR in a H3K79me2-independent manner.

To further dissect the mechanism through which DOT1L suppresses retroelements, we examined the potential interactors of DOT1L by IP-MS in HuH-7 cells (table S5). Among these high-confidence partners (fig. S5A), we validated the interaction between DOT1L and NPM1 (fig. S5B), since a recent study reported that Dot1l cooperates with Npm1 to inhibit the retrovirus MERVL expression in mouse embryonic stem cells (*27*). Importantly, depletion of NPM1 led to up-regulation of retroelements, accumulation of dsRNA, and induction of ISGs (fig. S5, C to F), similar to those of DOT1L deficiency. We next performed NPM1 ChIP-seq experiments and found that the binding pattern of NPM1 was similar to that of DOT1L in HuH-7 cells (fig. S5G). Around 66% of DOT1L peaks were overlapped with NPM1 in the genome wide (fig. S5, H and I). Notably, nearly 42.5% of the DOT1L-bound and -inhibited LINEs and LTR was also occupied by NPM1 (fig. S5, J to L), suggesting that DOT1L collaborates with NPM1 to repress these retroelements. Taken together, these data demonstrate that DOT1L-NPM1 complex suppresses retroelement expression to restrain cell-Intrinsic IFN response.

### DOT1L activates ZEB1 to repress the immune-related genes

DOT1L is typically associated with active genes via histone H3K79 methylation (*31*). Indeed, ∼40% of DOT1L peaks located at gene promoters and the corresponding histone H3K79me2 distributed across the gene bodies (Fig. 4J and fig. S6A). Of note, there were no detectable DOT1L binding or H3K79me2 signals at the *CD47* and *PD-L1* loci (fig. S6B), suggesting an indirect regulation. As expected, DOT1L loss diminished the levels of H3K79me2 at these genes in HuH-7 cells (fig. S6C). We reasoned that DOT1L depletion may also down-regulate the negative regulators to induce the IFN pathways. Hence, we integrated analysis of the H3K79me2 ChIP-seq and RNA-seq data, and found that ZEB1 appears to be a negative regulator of the IFN pathways (Fig. 5A), given that ZEB1 has been reported to promote immune escape in melanoma and lung cancer (*32, 33*). The expression of *ZEB1* was significantly decreased in DOT1L depleted HuH-7 cells by two independent sgRNAs, which was further validated by RT-qPCR and immunoblot analysis (Fig. 5, B to D). Moreover, treatment of HuH-7 cells with EPZ-5676 also decreased the mRNA levels of *ZEB1* (fig. S6D). Additionally, re-introduction of the WT DOT1L, rather than CI DOT1L, restored the *ZEB1* levels (fig. S6E). Consistent with that in HuH-7 cells, DOT1L deficiency or methyltransferase activity inhibition by EPZ-5676 also led to the reduction of *ZEB1* expression in HepG2 cells (fig. S6, F and G). Furthermore, the *DOT1L* expression is positively correlated with the *ZEB1* levels in liver according to the (Genotype-Tissue Expression) GTEx database (fig. S6H). Finally, according to the DOT1L fanChIP data, DOT1L directly occupied at the *ZEB1* promoter (Fig. 5E). DOT1L ablation in HuH-7 cells resulted in reduction of the histone H3K79me2 levels of the *ZEB1* gene body, H3K4me3 and H3K27ac levels, as well as the chromatin accessibility of the *ZEB1* promoter (Fig. 5E). These data together demonstrate that DOT1L directly contributes to the *ZEB1* activation.

**Fig. 5.**
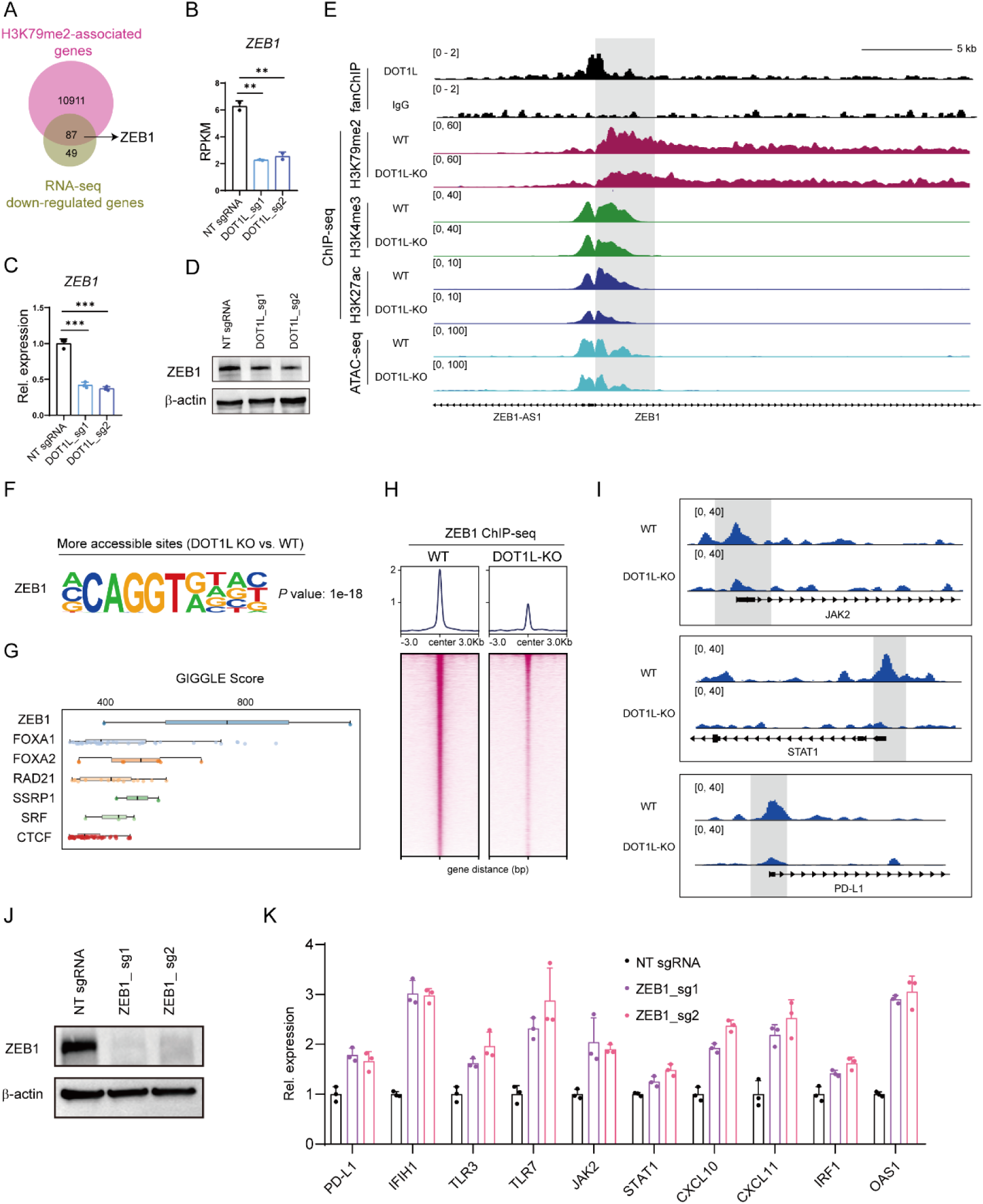
DOT1L activates ZEB1 to inhibit immune-related genes. (**A**) Venn diagrams of H3K79me2-associated genes and down-regulated genes in HuH-7 cells. (**B**) Expression levels of *ZEB1* by RNA-seq in WT and DOT1L-depleted HuH-7 cells. Results are shown as mean ± SD (n=2). \*\**P* < 0.01, unpaired Student’s *t*-test. RPKM, Reads Per Kilobase per Million mapped reads. (**C**) RT-qPCR analysis of *ZEB1* upon DOT1L depletion in HuH-7 cells. Results are shown as mean ± SD (n=3). \*\*\**P* < 0.001, unpaired Student’s *t*-test. (**D**) Immunoblotting analysis of ZEB1 protein levels using whole-cell lysates from WT and DOT1L KO HuH-7 cells. (**E**) DOT1L fanChIP tracks, H3K79me2, H3K4me3, and H3K27ac ChIP-seq signals, and ATAC-seq profiles of WT and DOT1L-depleted HuH-7 cells in the *ZEB1* locus. (**F**) Enrichment of ZEB1 motifs in the accessible chromatin regions specific to DOT1L-deficient HuH-7 cells by HOMER analysis. (**G**) Cistrome toolkit analysis of ATAC-seq data revealed that DNA-binding sites of ZEB1 were more open in the DOT1L-deficient HuH-7 cells. (**H**) Binding density heatmap of ZEB1 ChIP-seq signals in the DOT1L-deficient HuH-7 cells. (**I**) ChIP-seq tracks of ZEB1 in WT and DOT1L-depleted HuH-7 cells in the *JAK2*, *STAT1*, and *PD-L1* loci. (**J**) Immunoblotting analysis using whole-cell lysates from WT and ZEB1 KO HuH-7 cells. (**K**) Expression levels of immune-related genes by RT-qPCR analysis upon ZEB1 depletion in HuH-7 cells. Results are shown as mean ± SD (n=3).

Interestingly, the patients who had liver hepatocellular carcinoma (LIHC) with high *ZEB1* expression in tumors showed significantly reduced overall survival (fig. S6, I and J). Moreover, HOMER analysis and the Cistrome Data Browser (*34*) uncovered ZEB1 motif to be highly enriched in the peaks with increased chromatin accessibility upon DOT1L loss in HuH-7 cells (Fig. 5, F and G). More importantly, unbiased CRISPR screens targeting the human transcription factor library (unpublished data) showed that ZEB1 functioned as a potential repressor of PD-L1 in HuH-7 cells (fig. S6K). These data imply that ZEB1 contributes to the IFN pathway activation by DOT1L loss. To test our hypothesis, we performed ZEB1 ChIP-seq experiments and found that the global ZEB1 binding signals were remarkably reduced upon DOT1L depletion (Fig.5H and fig. S6L), consistent with the mRNA and protein levels of ZEB1 in DOT1L deficient HuH-7 cells (Fig. 5, B to D). The majority of ZEB1 binding peaks located at promoters (68%), and further motif analysis revealed that these binding peaks highly enriched binding sites for ZEB1 and STAT family (fig. S6, M and N), suggesting the role in immune response. In addition, ∼90% of the differential ZEB1 peaks sited at gene promoters (fig. S6O), indicating that ZEB1 directly modulates the target gene expression. Indeed, DOT1L loss remarkably weakened ZEB1 binding at the immune-related genes, including *PD-L1*, *JAK2*, *STAT1*, *IRF1*, and *IFIH1* loci in HuH-7 cells (Fig.5I and fig. S6P). Lastly, depletion of ZEB1 led to induction of the immune-related genes (Fig. 5, J and K). Of note, there was no detectable ZEB1 binding at the *CD47* locus and *ZEB1* depletion had no effect on its expression (fig. S6, Q and R). Meanwhile, ZEB1 loss showed minimal effect on the expression of TEs (fig. S6S). Collectively, these findings demonstrate that DOT1L promotes *ZEB1* expression to further repress the IFN pathway genes.

### DOT1L abrogation-induced IFN pathway activation sensitizes tumor response to ICB therapy

We next investigated biological consequences of Dot1l inhibition in murine HCC Hepa1-6 and Hep53.4 cells, as well as Lewis lung carcinoma (LLC) cell line. Dot1l ablation by CRISPR/Cas9 resulted in up-regulation of innate immune response and antiviral response pathways via GO enrichment analysis of the RNA-seq data, compared with WT Hepa1-6 cells (Fig. 6, A and B). GSEA revealed the up-regulation of IFN and JAK-STAT signaling pathways upon Dot1l depletion (Fig. 6C). Moreover, Dot1l loss also led to the up-regulation of retroelement expression and subsequent accumulation of dsRNA (Fig. 6, D and E). Consistently, treatment of Hepa1-6 cells with EPZ-5676 up-regulated the expression of immune-related genes (Fig. 6, F and G), despite that Hepa1-6 cells is less sensitive to EPZ-5676 than the human HuH-7 and HepG2 cell lines. In addition, depletion of Dot1l in Hep53.4 and LLC cells also led to accumulation of dsRNA and up-regulation of the immune-related gene expression. Again, treatment of Hep53.4 and LLC cells with EPZ-5676 increased the mRNA levels of immune-related genes (fig. S7, A to L). These data recapitulate our findings in human HuH-7 and HepG2 cells to some degree. Of note, Dot1l appeared not to activate *Zeb1* expression in Hepa1-6, Hep53.4, and LLC cells, which may be due to the evolutionary distinctiveness between humans and mice (fig. S7, M to O).

**Fig. 6.**
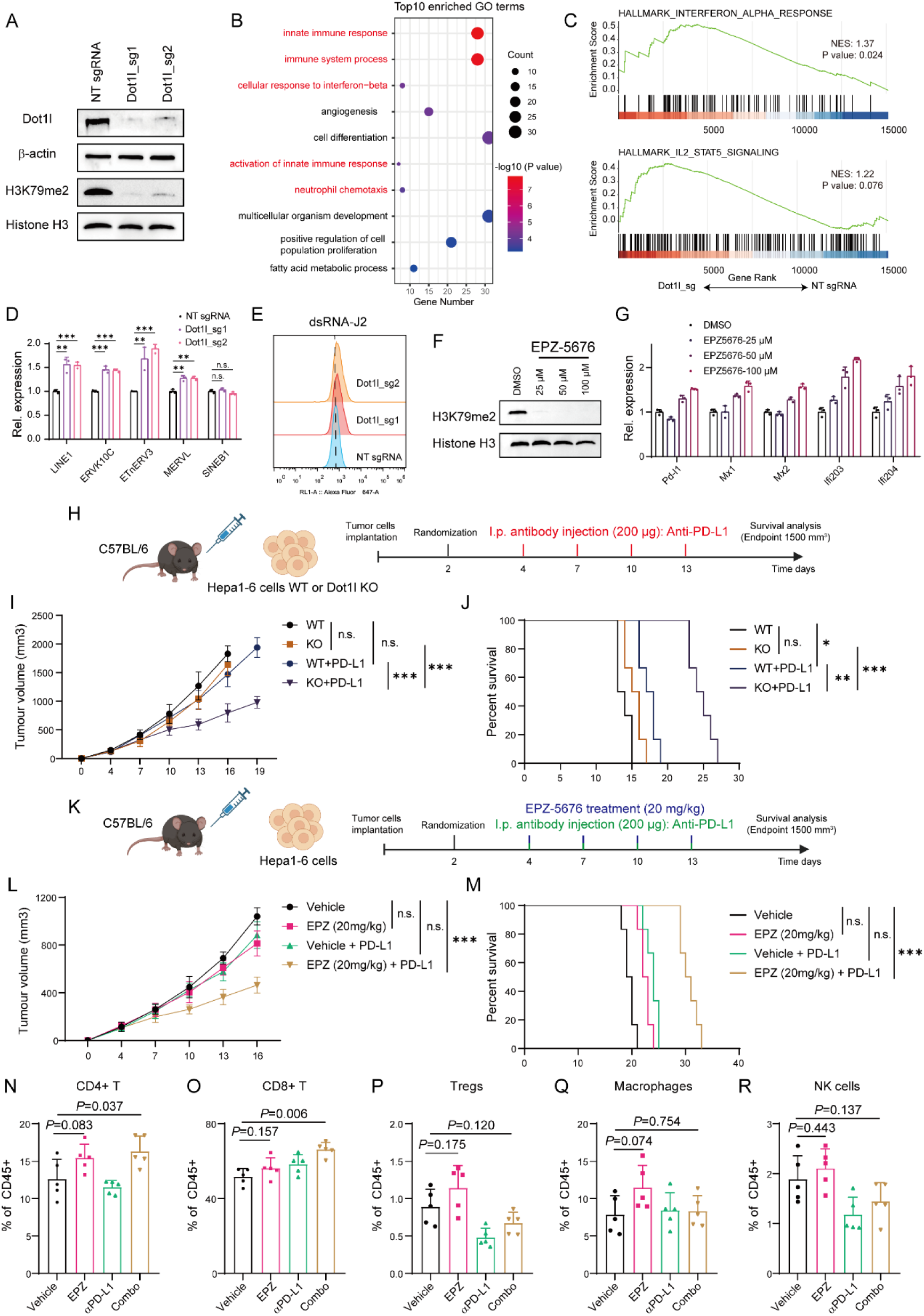
DOT1L deficiency or methyltransferase activity inhibition sensitizes tumors to ICB therapy. (**A**) Immunoblotting analysis using whole-cell lysates from WT and Dot1l KO Hepa1-6 cells. (**B**) GO analysis of RNA-seq data showing top 10 pathways that are up-regulated in Hepa1-6 cells upon Dot1l deficiency. (**C**) GSEA of RNA-seq data showing IFN (top) and IL2_STAT5 (bottom) gene sets that are up-regulated upon DOT1L depletion in Hepa1-6 cells. (**D**) RT-qPCR analyses of TEs in WT and Dot1l-depleted Hepa1-6 cells. Results are shown as mean ± SD (n=3). n.s., not significant, \*\**P* < 0.01, \*\*\**P* < 0.001, unpaired Student’s *t*-test. (**E**) Flow cytometry histograms of dsRNA staining of WT or Dot1l KO Hepa1-6 cells. (**F**) Immunoblotting analysis using whole-cell lysates from Hepa1-6 cells with EPZ-5676 treatment for 7 days. (**G**) RT-qPCR analysis of immune-related genes of Hepa1-6 cells with EPZ-5676 treatment for 7 days. Results are shown as mean ± SD (n=3). (**H**) Schematic of ICB treatments in mice injected with Hepa1-6 WT or Dot1l KO cells. (**I**) Analysis of tumor volume in C57BL/6 mice subcutaneously injected with Hepa1-6 WT or Dot1l KO cells reconstituted as in (H) and subsequently treated with isotype control or anti-PD-L1 antibodies (n=6). n.s., not significant, \*\*\**P* < 0.001, unpaired Student’s *t*-test. (**J**) Survival analysis conducted on the C57BL/6 mice described in (H). n.s., not significant, \**P* < 0.05, \*\**P* < 0.01, \*\*\**P* < 0.001, Logrank test. (**K**) Schematic of Dot1l inhibitor and ICB treatments in mice injected with Hepa1-6 cells. (**L**) Analysis of tumor volume in C57BL/6 mice subcutaneously injected with Hepa1-6 cells reconstituted as in (K) and subsequently treated with 20 mg/kg EPZ-5676 and anti-PD-L1 antibodies (n=6). n.s., not significant, \*\*\**P* < 0.001, unpaired Student’s *t*-test. (**M**) Survival analysis conducted on the C57BL/6 mice described in (L). n.s., not significant, \*\*\**P* < 0.001, Logrank test. (**N** to **R**), The intra-tumoral abundance of CD4+ T cells, CD8+ T cells, Tregs, NK, and Macrophages cells upon Dot1l inhibition alone or in combination (Combo) with anti-PD-L1 blockade. Results are shown as mean ± SD (n=5). *P* values were calculated by unpaired Student’s *t*-test.

Based on our in vitro findings, we asked whether Dot1l inhibition could trigger antitumor immunity in vivo. To this end, we used syngeneic mouse tumor models by subcutaneously injection of Hepa1-6 cells into C57BL/6 mice. However, depletion of Dot1l showed minimal effect on tumor growth (fig. S8A). In addition, drug treatment with the clinical stage inhibitor EPZ-5676 weakly affected the tumor size compared with the vehicle treatment (fig. S8B). Thus, we speculated that depletion of Dot1l or treatment with inhibitor alone was not sufficient to induce effective antitumor immunity. Previous study has shown that knockout of *Setdb1*, *Tasor* or *Mphosph8* sensitized LLC cells to anti-PD-1/CTLA-4, but had little effect on tumor growth without ICB (*12*). To test the hypothesis, we investigated whether Dot1l deficiency could enhance anti–PD-L1 inhibitor efficacy in the subcutaneous tumor model inoculated with Hepa1-6 cells (Fig. 6H). The results again showed little improvement by tumor growth curves and animal survival between WT and Dot1l KO groups, as well as WT treated with isotype control or anti–PD-L1 groups (Fig. 6, I and J). Importantly, Dot1l depletion dramatically enhanced the sensitivity of Hepa1-6 tumors to anti–PD-L1 therapy, resulting in markedly reduced tumor size and increased mouse survival (Fig. 6, I and J). Consistently, Hepa1-6 tumors in mice treated with combined EPZ-5676 and anti-PD-L1 therapy significantly decreased tumor size and thus increased animal survival (Fig. 6, K to M). To explore the impact on TME in these groups, we evaluated tumor-infiltrating immune cells by flow cytometry analysis (fig. S8C) and found a significant increase of immune cells including helper CD4+ and cytotoxic CD8+ T cells in the combinational group, compared with the vehicle-treated group (Fig. 6, N and O), whereas the amounts of regulatory T cells (Tregs), NK1.1 and macrophages did not change (Fig. 6, P to R). To validate our findings in a second in vivo model, we engrafted Hep53.4 WT or Dot1l KO cells into the flanks of recipient C57BL/6 mice. Similarly, Dot1l loss also enhanced the sensitivity of Hep53.4 tumors to anti– PD-L1 therapy, leading to reduced tumor size (fig. S8D). Again, Hep53.4 tumors with EPZ-5676 and anti-PD-L1 combination therapy strikingly reduced the tumor volume, compared with the control group (fig. S8E). Together, these results suggest that DOT1L loss can sensitize tumors to immunotherapy.

### DOT1L inhibition exhibits superior antitumor efficacy in humanized mice

Given that DOT1L has dual function in IFN pathway activation in humans, we generated the HuH-7 cell line-derived xenograft (CDX) in humanized immunocompetent mice (huHSC-NCG) to further explore the translational significance of our findings (Fig. 7A). The immune system of huHSC-NCG mice mainly reconstructed with human T cells and a large number of immature B cells, as well as a small number of NK cells. Of note, depletion of DOT1L or EPZ-5676 treatment did not influence the cell proliferation of HuH-7 in vitro (fig. S9, A and B). Importantly, treatment of huHSC-NCG mice with EPZ-5676 alone significantly decreased tumor volume and increased survival time (Fig. 7B and fig. S9C), displaying superior efficacy compared with that in murine tumors (discussed below). Flow cytometry revealed that the intra-tumoral infiltration of cytotoxic CD8+ T cells was significantly increased upon EPZ-5676 treatment (Fig. 7C). Of note, this treatment did not affect the body weight of mice (fig. S9D). These results show that DOT1L activity inhibition alone is sufficient to enhance antitumor immunity in humanized mouse tumor model.

**Fig. 7.**
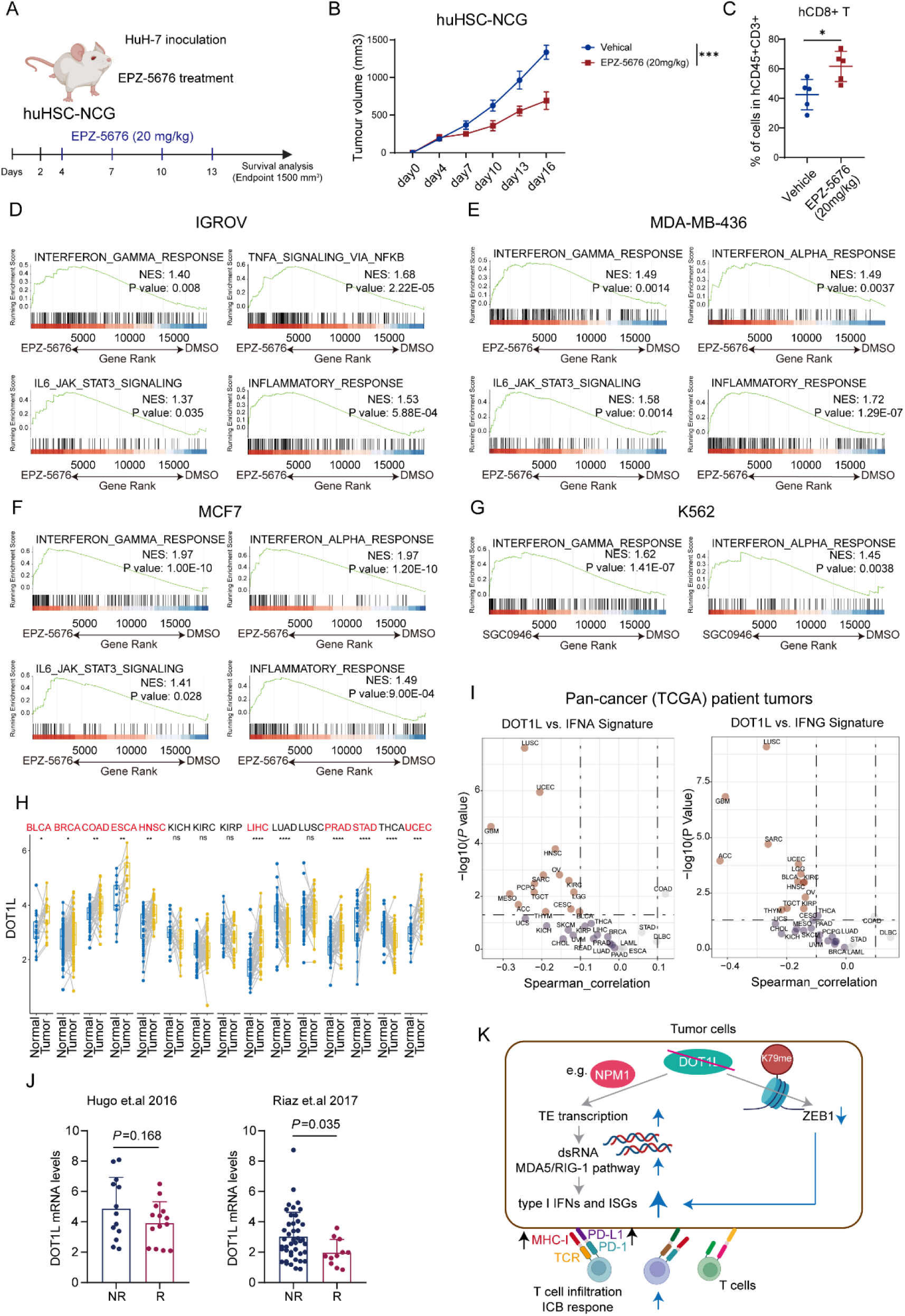
DOT1L inhibition enhances antitumor immunity in humanized mouse and DOT1L expression is conversely correlated with IFN signatures in multiple cancer cell lines and patient data sets. (**A**) Schematic of DOT1L inhibitor treatment in huHSC-NCG mice injected with HuH-7 cells. (**B**) Analysis of tumor volume in huHSC-NCG mice subcutaneously injected with HuH-7 cells reconstituted as in (A) and subsequently treated with 20 mg/kg EPZ-5676 (n=5). \*\*\**P* < 0.001, unpaired Student’s *t*-test. (**C**) The intra-tumoral abundance of hCD8+ T cells upon DOT1L inhibition treatment in huHSC-NCG mice (n=5). \**P* < 0.05, unpaired Student’s *t*-test. (**D** to **G**) GSEA of published RNA-seq data showing immune-related gene sets that are up-regulated upon inhibition the methyltransferase activity of DOT1L in multiple cancer cell lines. (**H**) Analysis of DOT1L expression in tumors and normal tissues from TCGA patients with the indicated cancer types. n.s., not significant, \**P* < 0.05, \*\**P* < 0.01, \*\*\**P* < 0.001, \*\*\*\**P* < 0.0001, wilcox.test. (**I**) Correlation analysis between DOT1L expression levels and z-scores of the IFN gene sets in different tumor types of TCGA. (**J**) DOT1L mRNA expression levels of responder and non-responder in ICB treatment clinical trials with anti-PD-1 treatment in melanoma. *P* values were calculated by unpaired Student’s *t*-test. (**K**) A proposed model for the dual function of DOT1L in regulating the immune response of tumors.

### DOT1L expression is negatively correlated with IFN signatures in multiple human cancer cell lines and patient data sets

We next interrogated published transcriptomic data from multiple cancer cell lines (*35–38*), treated with DOT1L inhibitor EPZ-5676 or SGC0946, to determine whether DOT1L generally represses the IFN pathway activation. GSEA revealed the up-regulation of IFN-dependent and cell-intrinsic immune signaling pathways in IGROV, MDA-MB-436, MCF7, and K562 cells (Fig. 7, D to G). To further explore the findings in human cancer patients, we first examined DOT1L mutations in the cBioPortal dataset and found that DOT1L was infrequently mutated, amplified, or deleted in human cancer (fig. S9E). Regardless, DOT1L mutation appeared to activate IFN pathway in multiple tumor types (fig. S9, F to H). Additionally, DOT1L was overexpressed in tumors compared with normal tissues in a number of cancer types based on the TCGA dataset (Fig. 7H). According to the DOT1L expression level, we divided patients of each cancer by DOT1L expression quartile (top 25%, high group and bottom 25%, low group) and compared overall survival and disease-free survival between the two groups. Our analysis showed that the DOT1L-high group had a significantly shorter overall survival and disease-free survival time than that of DOT1L-low group in multiple cancer types (fig. S9, I to K), suggesting that DOT1L overexpression is a poor prognostic factor. Further GSEA revealed that DOT1L expression inversely correlated with the gene signatures involved in IFN response in these tumor types (fig. S9, L to N), in agreement with the results from both human and mouse cell lines. Indeed, pan-cancer analysis exhibited a negative correlation between DOT1L expression and IFN signatures in a variety of tumor types based on the TCGA cancer patient datasets (Fig. 7I). Lastly, we analyzed two published RNA-seq datasets of pre-treatment cancer patient samples before anti-PD-1 therapy (*39, 40*) and found that responders to immunotherapy expressed lower levels of DOT1L than non-responders (Fig. 7J), in line with our finding that DOT1L loss sensitizes the response to ICB therapy. Taken together, these data suggest that DOT1L inhibition may generally improve cancer immunotherapy via inducing the IFN signaling pathways.

## DISCUSSION

In this study, through domain-focus CRISPR/Cas9 screening, we uncover the epigenetic regulator DOT1L functions as a repressor of innate immune signaling, and thus immune checkpoint *CD47* and *PD-L1* expression. Our study reveals that loss of DOT1L induces a dsRNA–MDA5/RIG-1–type I IFN response axis that causes tumor cell–intrinsic antiviral response activation in an enzymatic-dependent manner. Moreover, we demonstrate that DOT1L acts as a dual regulator of TEs and *ZEB1* expression that promotes tumor immune evasion in human HCC cell lines (Fig. 7K). Furthermore, DOT1L depletion or clinical stage inhibitor EPZ-5676 treatment greatly improves ICB therapy outcomes in murine tumors. Particularly, EPZ-5676 treatment alone in humanized mice effectively enhance antitumor immunity. These findings thus shed important insight into DOT1L in regulating tumor immunity and propose the translational potential of DOT1L inhibitor as a strategy to enhance response to checkpoint immunotherapy for cancer patients.

DOT1L preferentially occupies the proximal transcribed region of active genes in diverse mammalian cell types (*25*). In contrast, several evidences revealed that DOT1L is also associated with transcriptional repression (*27, 28, 41–43*). Indeed, we show that DOT1L directly inhibits the chromatin accessibility and transcription of TEs partially through interacting with NPM1 (fig. S5), in line with a recent study that the Dot1l-Npm1 complex repressed the retrovirus MERVL in mouse embryonic stem cells (*27*), suggesting that DOT1L noncanonically functions as a transcription repressor for a subset of TEs including LINEs and LTR. However, the remaining up-regulated retroelements seem not to be directly regulated by DOT1L-NPM1 complex (fig. S5J). It will be intriguing to characterize more potential factors mediating DOT1L-inhibited TEs in the future, as well as associated transcription factors.

A previous literature has shown that DOT1L modulates cell-fate determination and transcriptional elongation independent of its catalytic activity (*44*). We demonstrate that DOT1L represses the antitumor immunity in an enzymatic-dependent manner. Currently, DOT1L is best known for its ability to catalyze the histone substrate H3K79me2 which is associated with active gene expression (*25, 26, 45, 46*). However, according to our data, despite DOT1L bound at the promoters of ∼6000 genes that enriched histone H3K79me2 across the gene bodies in HuH-7 cells (fig. S5A), only 87 genes were significantly down-regulated upon DOT1L loss (Fig. 5A). These findings indicate that DOT1L activating gene expression likely relies on specific epigenetic contexts, rather than singular H3K79me2 marks. Indeed, a latest study reported that DOT1L exhibits opposite effects on gene expression depending on chromatin environment during mouse spermatogenesis (*28*). Regardless, we demonstrate that *ZEB1* is one of the DOT1L targets involved in H3K79me2 modifications (Fig. 5). In contrast, compared with the active gene promoters, these DOT1L-inhibited TE regions enriched background levels of H3K79me2 that were inert in response to DOT1L depletion (fig.S4M and S5A), arguing against H3K79me2-dependent repression of the TEs. Meanwhile, DOT1L has been reported to methylate non-histone substrate, such as androgen receptor and RAP80 (*47, 48*), thus we cannot exclude the possibility that DOT1L represses the expression of TEs via unknown non-histone substrates.

Epigenetic repression of immunogenicity has become a major focus, by which tumors evade immune system eradication and become resistant to immunotherapies. Increasing evidences have revealed that the epigenetic regulators are the repressors of TEs, and thus cell-intrinsic innate immunity (*10, 11, 13, 14, 22, 23, 49–51*). We demonstrate that DOT1L loss-induced TE expression and resulting IFN signaling activation are conserved between humans and mice during evolution. Moreover, DOT1L activates *ZEB1* expression to further promote immune evasion in humans. However, *Zeb1* appeared not to be regulated by Dot1l in mice, exhibiting species-specific gene regulation (*52, 53*). Consistently, a previous study has shown the differences of transcriptional landscapes between human and mouse tissues, suggesting that gene expression is more similar among tissues within the same species than between corresponding tissues of two species (*54*). Additionally, Dot1l deficiency or methyltransferase activity inhibition alone was insufficient for tumor killing, which may be partially due to the absence of the Dot1l-Zeb1 axis in murine tumors. Nevertheless, combination EPZ-5676 with anti-PD-L1 therapy reprograms the TME in murine tumor models, increasing accumulation of CD4+ and CD8+ T cells, similar to that of combination strategies with EZH2 inhibitors and anti-PD-1 immunotherapy in prostate cancer (*22*). More importantly, treatment of humanized mice with EPZ-5676 alone can enhance antitumor immunity and thus reduce tumor size. Moreover, we found an inverse correlation between DOT1L expression and IFN signatures in human cancer cell lines and cancer patients, suggesting that DOT1L expression may be an informative biomarker in a variety of cancer types.

In sum, our findings highlight the therapeutic potential of targeting DOT1L or utilizing the clinical stage inhibitor EPZ-5676 in combination with PD-(L)1 blockade for HCC treatment, which may be a generally applicable new strategy for cancer immunotherapy.

## MATERIALS AND METHODS

### Study design

This study aimed to uncover novel epigenetic factors modulating the expression of immune checkpoints or tumor cell-intrinsic immunity. We carried out CRISPR genetic screens targeting epigenetic factors in hepatocellular carcinoma cell line and identified DOT1L as a potential suppressor of tumor cell-intrinsic immunity. For animal experiments, at least five mice per group were used with random assignment. Investigators were blinded to animal study groups and treatments. Animal experimental end points were predetermined at the beginning of the study. All in vitro experiments were performed at least three biological replicates and appropriate statistical tests were performed. For RNA-seq, ChIP-seq and ATAC-seq were performed with two biological replicates, and no data were excluded.

### Cell culture

HEK293T cells (RRID:CVCL_0063), HuH-7 cells (RRID:CVCL_0336), HepG2 cells (RRID:CVCL_0027), Hepa1-6 cells (RRID:CVCL_0327), Hep53.4 cells (RRID:CVCL_5765), and LLC cells (RRID:CVCL_4358) were maintained in DMEM media (Gibco) with 10% fetal bovine serum and 1% penicillin-streptomycin solution. For inhibitor treatment, cells were treated with DOT1L inhibitor EPZ-5676 (MedChemExpress, HY-15593) for 7 days. All cells were routinely tested for *Mycoplasma* and found to be free of contamination. All cells were cultured at 37°C with 5% (v/v) of CO_2_.

### Mice

C57BL/6J male mice and huHSC-NCG female mice were purchased from GemPharmatech. C57BL/6J male mice were six to eight weeks of age and huHSC-NCG female mice were 16 weeks of age at the beginning of tumor studies. Animals were housed in specific-pathogen-free facilities at the Department of Laboratory Animal Science of Fudan University. All animal studies and protocols were performed in accordance with a protocol approved by the Institutional Animal Care and Use Committee of Fudan University.

### Plasmid construction

To prepare sgRNA plasmids for CRISPR/Cas9 systems, complementary oligonucleotide pairs with 5’ overhangs of CACC and AAAC were annealed to generate dsDNA and then sub-cloned into the BsmbI-digested lentiviral sgRNA-EFS-GFP/mCherry vector (Addgene #65656). The primers are listed in table S6.

To prepare shRNA plasmids for gene knockdown, complementary oligonucleotide pairs with 5’ overhangs of CCGG and TTTTTG were annealed and then sub-cloned into the AgeI and EcoRI-digested pLKO.1 vector (Addgene #10878). The primers are listed in table S6.

For the generation of DOT1L overexpression cell lines, the DOT1L cDNA with HA or FLAG tag was constructed into the XbaI and BamHI-digested lenti-Cas9-Blast (LRG, Addgene #62962) vector.

### Lentiviral transduction

Lentivirus was prepared by transfecting HEK293T cells for each stable cell line construction. Briefly, about 10 μg lentivirus transfer plasmid, 7.5 μg packaging helper plasmid (psPAX2, Addgene #12260), 5 μg envelope helper plasmid (pMD2.G, Addgene #12259) and 80 μl of 1mg/ml polyethylenimine (PEI) were mixed and incubated for 15 mins. The mixture was added to HEK293T cells grown in 10 cm plates to 90% confluence. After 8 h, the medium was replaced with fresh complete medium. The virus was collected after transfection for 48 h. For adherent cells infection, the mixture of virus, medium, 8 μg/ml polybrene and 10 mM HEPES was added and cultured for 24 h. After that, the medium was replaced with fresh complete medium. The transfected cells were selected in 1 μg/ml puromycin for 7 days or 10 μg/ml blasticidin for 14 days.

### CRISPR sgRNA library generation and screen

The sgRNAs library targeting human chromatin regulators and proteins containing bromodomain were performed as described previously (*19*). Briefly, HuH-7-Cas9 cells were transduced with the library at a low multiplicity of infection (MOI ∼0.3). About 5 × 10^6^ cells were infected in total to yield 1000 × coverage of the sgRNA library in the GFP+ population and sorted by GFP+ Fluorescence activated Cell Sorting (FACS) after 48 h. Transduced cells were cultured in DMEM media for an additional 7 days. After that, the cells were divided into two groups with or without IFN treatment for 48 h. On day 11 post-infection, cells were stained for CD47/PD-L1, and sorted into CD47/PD-L1 high and CD47/PD-L1 low populations. The genomic DNA was extracted from these samples with phenol-chloroform, followed by ethanol precipitation. The sgRNAs library amplificon was used with Phanta Flash Master Mix (Vazyme). After that, the PCR products were pooled and column purified with QIAGEN PCR purification kit, subjected to Illumina library construction and sequencing. MAGeCK software (*55*) was used for sgRNA library analysis. Normalized read counts of sgRNAs in CD47/PD-L1 high and CD47/PD-L1 low populations were log2 transformed in RStudio software. All sgRNA sequences used in this study and screen results are provided in table S1.

### RNA extraction and RT-qPCR

Total RNA was extracted using the FastPure Cell/Tissue Total RNA Isolation Kit V2 (Vazyme). According to the manufactureŕs instructions, about 1 μg of total RNA was used in reverse transcription (Vazyme). The cDNA template was diluted and used for qPCR with SYBR Green master mix (Roche). All qPCR experiments were performed with at least three biological replicates. Data statistical tests were performed with GraphPad Prism (v8.0). Primers for qPCR are listed in table S6.

### Flow Cytometry

For detecting proteins on the cell membrane, collected cells were washed with PBS and stained with anti-CD47 (BioLegend, 323123), anti–PD-L1 (BioLegend, 329708), and anti–HLA-A, B, C (BioLegend, 311410) antibodies diluted with PBS. The fluorescence levels were compared with isotype control antibodies.

For intracellular flow cytometry, collected cells were fixed with 4% formaldehyde in PBS and permeabilized with 0.1% Triton X-100 in PBS. The cells were then incubated with primary antibody against dsRNA (J2; SCICONS, no. 10010200) or isotype controls antibody for 1 h. After that, the secondary goat anti-mouse IgG (H+L) conjugated with Alexa Fluor 647 (Thermo Fisher Scientific, A32728) was added to the cells and incubated for 30 mins.

Single-cell suspensions of tumors were prepared using a gentleMACS tissue dissociation system. The cells were incubated with primary antibody against BV510-mCD45 (BioLegend, 103137), PE-cy7-mCD3 (Thermo Fisher Scientific, 25-0031-82), PerCP-Cy5.5-mCD8 (BioLegend, 100734), FITC-mCD4 (Thermo Fisher Scientific, 11-0042-85), BV421-mCD25 (BioLegend, 102043), PE-mFoxp3 (Thermo Fisher Scientific, 12-5773-82), BV786-mCD11b (Thermo Fisher Scientific, 417-0112-82), BUV495-mF4/80 (BD Biosciences, 565614), APC-mNK1.1 (BioLegend, 108710) for staining. For humanized mice, the cells were incubated with primary antibody against BUV395-hCD45 (BD Biosciences, 563792), BV421-hCD3 (BD Biosciences, 563798), APC-hCD4 (BioLegend, 300514), and PE-Cy7-hCD8 (Thermo Fisher Scientific, 25-0086-42) for staining. After staining, cells were centrifuged at 600 × g for 3-5 mins, and washed twice before being analyzed or sorted. All stained cells were analyzed on an Attune™ NxT Flow Cytometer. Data were analyzed using the FlowJo software.

### Animal experiments

About 5 × 10^6^ WT and Dot1l-KO Hepa1-6 or Hep53.4 cells were resuspended in PBS and injected subcutaneously into the flank of recipient C57BL/6 mice. For xenograft tumor modeling in humanized mice, about 5 × 10^6^ HuH-7 cells in 50 mL ice-cold PBS mixed with 50 mL Matrigel (Corning, 356234) were injected subcutaneously into the flank of recipient huHSC-NCG mice. For Dot1l inhibitor treatment, C57BL/6 and huHSC-NCG mice were treated with 20 mg/kg EPZ-5676 twice per week for a total of 2 weeks after tumor engraftment on day 4. For immunotherapy treatment, anti-PD-L1 antibodies (BE0101) were purchased from Bio X cell. C57BL/6 mice were treated with 200 µg anti-PD-L1 antibodies or isotype control on day 4 twice a week after tumor injection and continuing four times. All tumor volumes were measured by digital calipers and calculated using the formula: (width * width * length)/2. Tumor growth and survival data were plotted in GraphPad Prism (v8.0).

### Immunoblot analysis

Protein was extracted from about 1-2 × 10^6^ cells with the RIPA lysis buffer (Thermo Fisher Scientific, 89900) with 1 × protease inhibitors (Roche) and 1 × phenylmethylsulfonyl fluoride (PMSF) for 30 mins on ice and then quantified by Pierce BCA Protein Assay Kit (Thermo Fisher Scientific, 23225). Protein lysate was denatured at 95°C for 5-10 mins and then about 20-30 μg proteins were separated by SDS-PAGE and transferred to nitrocellulose membranes. Transferred protein was immunoblotted using primary antibodies against DOT1L (Cell Signaling Technology, 90878), H3K79me2 (Abcam, ab3594), H3 (Proteintech, 17168-1-AP), HRP-conjugated-β-actin (Proteintech, HRP-60008), HA-tag (Cell Signaling Technology, 3724), NPM1 (Proteintech, 10306-1-AP), and ZEB1 (Proteintech, 21544-1-AP). The secondary HRP-conjugated Goat Anti-Rabbit IgG(H+L) was purchased from Proteintech (SA00001-2). The membranes scanned by the ChemiDoc Imaging System (Bio-Rad).

### IP-MS/IP-WB

About 5-10 × 10^8^ HuH-7 cells were washed with ice cold PBS twice, lysed with lysis buffer A (10 mM HEPES pH 7.5, 1.5 mM MgCl_2_, 10 mM KCl, 1 mM DTT, and 1 mM PMSF) for 10 mins on ice and then resuspended with nuclei lysis buffer C (20 mM Tris-HCl pH 7.9, 25% glycerol, 420 mM NaCl, 1.5 mM MgCl_2_, 0.1% NP-40, 0.2 mM EDTA, 1 mM DTT, and 1 mM PMSF). After that, the nuclei were treated with Benzonase nuclease on ice for 30 mins and harvested by scraping. The soluble nuclear proteins were then separated from the insoluble chromatin fraction by centrifugation at 45,000 rpm for 1.5 h and then incubated with anti-FLAG beads (Thermo Fisher Scientific) with rotations overnight at 4°C. On the second day, after one wash with buffer 150 (20 mM Tris-HCl pH 7.9, 25% glycerol, 150 mM NaCl, 1.5 mM MgCl_2_, 0.1% NP-40, 0.2 mM EDTA, 1 mM DTT, and 1 mM PMSF), one wash with buffer 350 (20 mM Tris-HCl pH 7.9, 25% glycerol, 350 mM NaCl, 1.5 mM MgCl_2_, 0.1% NP-40, 0.2 mM EDTA, 1 mM DTT, and 1 mM PMSF), and two washes with buffer 150, the beads were eluted with reducing SDS-loading buffer, heated at 95°C for 10 mins, and analyzed by SDS–PAGE for subsequent Nano-HPLC-MS/MS analysis.

For endogenous DOT1L immunoprecipitation, the nuclei lysate was mixed with 10 μg of primary antibody (Bethyl Laboratories, A300-954A) or isotype control IgG (abcam, ab172730) and incubated with rotations overnight at 4°C. The second day, the protein A/G beads were added and incubated with 3 h and washed in wash buffer 150 and 350, followed by eluting with reducing SDS-loading buffer, heating at 95°C for 10 mins, and analyzing by SDS–PAGE.

### Nano-HPLC-MS/MS analysis

Trypsin (12.5 ng/μl) was used for extract proteins in the gel slices. The peptides in the supernatant were collected after extract twice with solution (5% formic acid in 50% acetonitrile). Then, the tryptic peptides were subjected to NSI source, followed by tandem mass spectrometry (MS/MS) in Orbitrap Exploris 480 MS coupled online to the UPLC. After that, tandem mass spectra were extracted by Proteome Discoverer software (Thermo Fisher Scientific, version 3.0) and searched against Human database assuming the digestion enzyme trypsin. The acceptance criteria for identifications was the false discovery rate (FDR) should be less than 1% for peptides and proteins. Proteins identified in both replicate samples were subjected to downstream analysis.

### Total RNA-seq or RNA-seq

Total RNA was extracted as described above. Ribosomal RNA was removed by Ribo-off rRNA Depletion Kit (Human/Mouse/Rat) according to the manufacturés protocol (Vazyme) for total RNA-seq. RNA-seq experiments were performed using VAHTS Universal V6 RNA-seq Library Prep Kit for Illumina according to the manufacturés protocol (Vazyme). All total RNA-seq or RNA-seq libraries were performed with two biological replicates and sequenced on an Illumina NovaSeq 6000 platform.

### Chromatin immunoprecipitation (ChIP)

About 2 × 10^7^ HuH-7 cells were cross-linked with 1% formaldehyde at room temperature for 10 mins, followed by quenching with 0.1375 M glycine for 5 mins. Fixed cells were washed twice with ice cold PBS and resuspended in the cell lysis buffer (10 mM Tris pH 8, 10 mM NaCl, 0.2% Igepal) and nuclei lysis buffer (50 mM Tris pH 8, 1% SDS, 10 mM EDTA) with 1 × PMSF and 1 × protease inhibitors for 10 mins with rotations at 4°C. The nuclei were sonicated for 30 mins at 4°C (a train of 30 s on and 30 s off for 30 cycles). After removal of the insoluble debris, the cell lysate was pre-cleared with 50 μl Protein A/G agarose beads for 2 h at 4°C. After that, the supernatant was added Protein A/G beads pre-bound with 3-5 μg of specific antibodies against ZEB1 (Proteintech, 21544-1-AP), H3K27ac (Abcam, ab4729), H3K4me3 (Abcam, ab8580), and H3K79me2 (Abcam, ab3594) overnight. On the second day, after one wash with buffer I (20 mM Tris pH 8, 50 mM NaCl, 2 mM EDTA, 1% Triton X-100, 0.1% SDS), two washes with high salt buffer (20 mM Tris pH 8, 0.5 M NaCl, 2 mM EDTA, 1% Triton X-100, 0.01% SDS), one wash with buffer II (10 mM Tris pH 8, 0.25 M LiCl, 1 mM EDTA, 1% Igepal, 1% sodium deoxycholate), and two washes with TE (10 mM Tris pH 8, 1 mM EDTA pH 8), the beads were eluted twice with 100 μl elution buffer. And then, NaCl and RNase A were added to samples and incubated at 37°C for 30 mins. After that, proteinase K was added to samples and incubated at 65°C overnight. The DNA was purified with phenol-chloroform, followed by ethanol precipitation for library preparation.

For NPM1 (Proteintech, 10306-1-AP) ChIP experiments, the same protocol was used with one adaptation. Cells were first crosslinked in 10 ml PBS with 2 mM EGS for 20 mins at room temperature, followed by addition of 1% formaldehyde with crosslinking for 10 mins at room temperature. The DNA library was constructed according to the instructions of VAHTS Universal DNA Library Prep Kit for Illumina (Vazyme). All ChIP-seq libraries were sequenced on an Illumina NovaSeq 6000 platform. All ChIP experiments were performed with two biological replicates.

### fanChIP

Fractionation-assisted native ChIP (fanChIP) experiments were described as previously with modifications (*29*). About 1 × 10^7^ HuH-7 cells were washed once with ice cold PBS and resuspended in the CSK buffer (10mM PIPES pH 6.8, 100 mM NaCl, 3 mM MgCl_2_, 1mM EGTA pH 7.6, 0.3 M Sucrose, 0.5% Triton X-100, 0.5 mM DTT, and 0.5 mM Sodium butyrate) for 5 mins on ice. After that, the cell lysate was resuspended with MNase buffer (50mM Tris-HCl pH 7.5, 4 mM MgCl_2_, 1mM CaCl_2_, 0.3 M Sucrose, 0.5 mM DTT, and 0.5 mM Sodium butyrate) and incubated with 0.5 unit of MNase (Sigma-Aldrich) at 37°C for 5 mins, followed by quenching with by 0.5 M pH 8.0 EDTA. The cell lysate was resuspended with lysis buffer (20mM Sodium phosphate pH 7.0, 30 mM Sodium pyrophosphate 10·H_2_O, 250 mM NaCl, 10 mM NaF, 5 mM NaF EDTA pH 8.0, 10% Glycerol, and 0.5 mM DTT) for 5 mins on ice. After that, the supernatant was added 1 μg specific antibodies against DOT1L (Cell Signaling Technology, 77087) or isotype control IgG (Abcam, ab172730) with rotations overnight at 4°C. On the second day, the magnet beads (Thermo Fisher Scientific) were added, incubated for 3 h, and then washed five times with fanChIP buffer at 4°C. The DNA was eluted with elution buffer (50 mM NaHCO_3_ and 1% SDS) and purified with phenol-chloroform, followed by ethanol precipitation for library preparation.

### ATAC-seq

ATAC-seq experiments were performed using Hyperactive ATAC-Seq Library Prep Kit for Illumina according to the manufacturer’s instructions (Vazyme). All ATAC-seq libraries were sequenced on an Illumina NovaSeq 6000 platform. All ATAC-seq experiments were performed with two biological replicates.

### Tatol RNA-seq or RNA-seq data processing and analysis

For Tatol RNA-seq or RNA-seq experiments, raw FASTQ files were aligned to the human (GRCh38/hg38) or mouse reference genome (GRCm38/mm10) using HISAT2 (*56*) with default parameters. SAMtools (*57*) was applied to transform SAM into the BAM format. FeatureCounts (*58*) was used to calculate the read counts of each transcript. TEtranscripts (*59*) was used to calculate the read counts of transposable elements. Analyzing repeat expression on repetitive elements was calculated by Homer (*60*) analyseRepeats.pl.

For differential expression analysis, we used *P* < 0.05 and fold-change > 1.5 as thresholds to identify differentially expressed genes by DESeq2 package (*61*). For differential transposable elements analysis, we used *P* < 0.05 and fold-change > 1 as thresholds. Commonly changed genes in both independent sgRNAs were considered to be significant. DAVID (*62*) was used for the GO and KEGG enrichment analysis. ClusterProfiler package (*63*) was used for Gene Set Enrichment Analysis. Data statistical tests were performed with GraphPad Prism.

### ChIP-seq and fanChIP data processing and analysis

For ChIP-seq and fanChIP experiments, raw FASTQ files were aligned to the human reference genome (GRCh38/hg38) using Bowtie 2 (*64*). SAMtools (*57*) was applied to transform SAM into the BAM format and removed PCR duplicates. Bigwig files were generated using deepTools (*65*) and the signal was normalized using the counts per million (CPM), followed by visualizing in the Integrative Genomics Viewer (IGV).

Peaks were generated by MACS2 (*66*) with the default parameters. The final peaks were those overlapped by both biological replicates, created by BEDTools (*67*). Peak annotation and sequence motif analysis were carried out with Homer (*60*). For differential peak analysis, we used *P* < 0.05 and fold-change > 1 as thresholds by DESeq2 package (*61*). Heatmaps were generated by deeptools (*65*) computeMatrix and plotHeatmap.

### ATAC-seq data processing and analysis

For ATAC-seq experiments, raw FASTQ files were aligned to the human reference genome (GRCh38/hg38) using Bowtie 2 (*64*). All unmapped reads and PCR duplicates were removed by SAMtools (*57*). Bigwig files were generated and normalized using the CPM options in deepTools (*65*). MACS2 (*66*) was used to call peak with the default parameters. Peak annotation and sequence motif analysis were carried out by Homer (*60*). Differential ATAC peaks were analyzed by DESeq2 package (*61*) with *P* < 0.05 and fold-change > 1 as thresholds. Heatmaps were generated by deeptools (*65*) computeMatrix and plotHeatmap.

### Statistical analysis

Unpaired Student’s t test was used to analyze the differences between two groups. Survival curves were compared using the logrank test. The DOT1L expression levels between normal tissues and tumors from TCGA patients were compared with wilcox.test. Results were presented as the mean ± SD. *P* values with GraphPad Prism and R. *P* ≤ 0.05 was represented as ‘*’, *P* ≤ 0.01 was represented as ‘**’ and *P* ≤ 0.001 was represented as ‘***’, respectively.

## Supplementary Materials

**The PDF file includes:** Figs. S1 to S9

## Other Supplementary Material for this manuscript includes the following

Tables S1 to S6

## Acknowledgments

We thank all members of our laboratory for helpful comments and discussions. We also acknowledge the Medical Science Data Center at Shanghai Medical College of Fudan for providing the data analysis platform.

## Author contributions

S.X. and X.L. conceived the study and designed the experiments. S.X. carried out most the cell-based experiments. R. G. performed the CRISPR screen. S.X. and R. G. conducted mouse studies. S. L. analyzed the TCGA database. J. W., Y. S., C. P., and Q. F. helped with the experiments. S.X. analyzed RNA-seq, ChIP-seq, and ATAC-seq data. M. L., F. L., and J. F. provided the suggestion for experiment design and manuscript writing. J. C. and X.L acquired funding. S.X., J. C., and X.L. wrote the manuscript with input from all authors.

## Data and materials availability

The data generated in this study are publicly available in Gene Expression Omnibus (GEO).

